# Potent neutralizing antibodies in the sera of convalescent COVID-19 patients are directed against conserved linear epitopes on the SARS-CoV-2 spike protein

**DOI:** 10.1101/2020.03.30.015461

**Authors:** Chek Meng Poh, Guillaume Carissimo, Bei Wang, Siti Naqiah Amrun, Cheryl Yi-Pin Lee, Rhonda Sin-Ling Chee, Nicholas Kim-Wah Yeo, Wen-Hsin Lee, Yee-Sin Leo, Mark I-Cheng Chen, Seow-Yen Tan, Louis Yi Ann Chai, Shirin Kalimuddin, Siew-Yee Thien, Barnaby Edward Young, David C. Lye, Cheng-I Wang, Laurent Renia, Lisa F.P. Ng

## Abstract

The ongoing SARS-CoV-2 pandemic demands rapid identification of immunogenic targets for the design of efficient vaccines and serological detection tools. In this report, using pools of overlapping linear peptides and functional assays, we present two immunodominant regions on the spike glycoprotein that were highly recognized by neutralizing antibodies in the sera of COVID-19 convalescent patients. One is highly specific to SARS-CoV-2, and the other is a potential pan-coronavirus target.

## Main

In December 2019, a cluster of pneumonia cases of unknown etiology was reported in the city of Wuhan in the province of Hubei. The previously unidentified pathogen, which induces symptoms resembling an infection by the Severe Acute Respiratory Syndrome Coronavirus (SARS-CoV), was later identified as a novel coronavirus, SARS-CoV-2 [1]. Within a span of four months, there are more than 750,000 laboratory-confirmed cases of human Coronavirus Disease 2019 (COVID-19), with over 35,000 deaths across 199 countries and territories (For up to date information consult https://www.who.int/emergencies/diseases/novel-coronavirus-2019/situation-reports/). After being declared a pandemic by World Health Organization (WHO) on 11^th^ March 2020, there is a compelling need to understand and develop effective therapeutic interventions against SARS-CoV-2.

SARS-CoV-2 uses the spike (S) glycoprotein to bind to the angiotensin-converting enzyme 2 (ACE2) receptor with a better affinity than SARS-CoV S glycoprotein for entry [2]. Thus, blocking the binding to ACE2, or blocking host protease cleavage to release the fusion peptide is an efficient strategy to prevent coronavirus entry [3-5]. To date, one study has assessed the immunogenicity of structural domains of recombinant SARS-CoV-2 S protein [6]. At the time of writing, findings on SARS-CoV-2 linear epitopes remain limited to bioinformatics prediction of human B and T-cell epitopes using SARS-CoV as a model [7-9]. Five regions on the S glycoprotein of SARS-CoV (residues 274-306, 510-586, 587-628, 784-803 and 870-893) were predicted to be associated with a robust immune response [7], while other studies reported candidate epitopes [8, 9] that require validation with human patient samples.

In this brief communication, we report the antibody profiles of COVID-19 patients, and the identification of two immunodominant linear B-cell epitopes on the S glycoprotein of SARS-CoV-2 that are crucial in controlling infection. A total of 25 convalescence serum samples collected during the current COVID-19 outbreak in Singapore were screened at 1:1000 dilution for neutralizing antibodies against a pseudotyped lentivirus expressing SARS-CoV-2 S glycoprotein tagged with a luciferase reporter (Figure 1a). Of the 25 patients tested, six patients (2, 4, 6, 7, 11, 20) with sufficient amount of serum samples that displayed a good neutralizing activity were selected for further functional characterization. Sera from all patients showed similar IC_50_, ranging from a titre of 694 to 836, except patient 20, who showed the strongest neutralizing activity with an IC_50_ of 1603 (Figure 1b).

**Figure 1.**
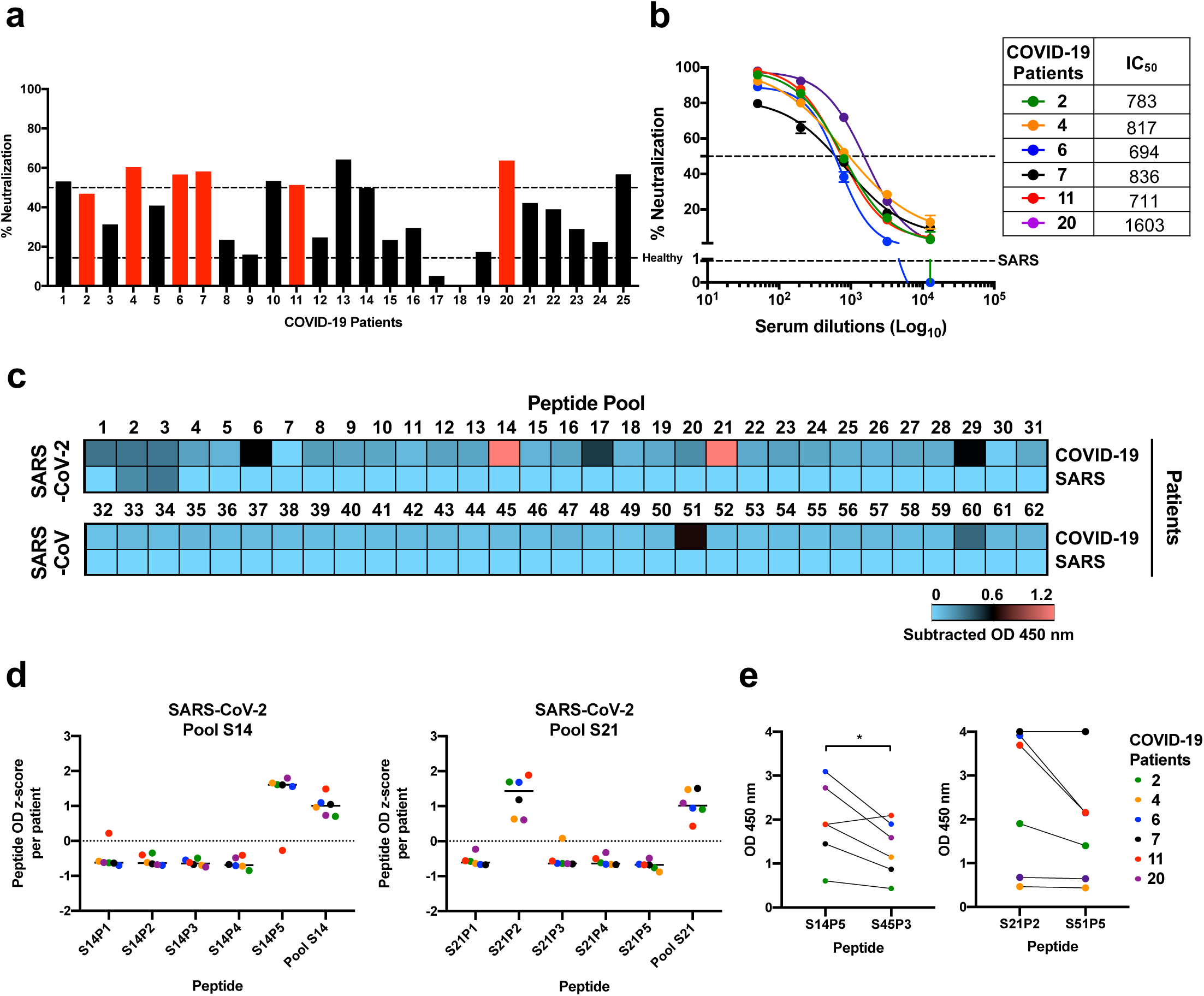
COVID-19 patients elicit neutralizing and specific antibody response against SARS-CoV-2. **(a-b)** Sera of COVID-19 patients (n=25) were mixed with pseudotyped lentiviruses expressing SARS-CoV-2 spike (S) glycoprotein protein, prior to incubation with CHO-ACE2 cells for 48 h. Infection levels were determined by luciferase assay, and percentage of infected cells were analyzed. Healthy control and recovered SARS patients (n=5) were also conducted in parallel. **(a)** Percentage of virus neutralization at 1:1000 sera dilution. IC_50_ titer is the reciprocal of sera dilution at which half-maximal neutralization was observed (dotted lines). **(b)** Dose-response neutralization kinetics from sera (1:50 to 1:12800 dilutions) of selected COVID-19 patients (n=6). Dotted lines indicate 50% neutralization or percentage neutralization of SARS patients (n=5) at 1:100 sera dilution. **(c)** Preliminary mapping of linear B cell epitopes within SARS-CoV-2 and SARS-CoV S protein. Sera of COVID-19 (n=6) and SARS patients (n=5) at 1:1000 dilution were subjected to peptide-based ELISA for IgG using peptide pools of the S protein of SARS-CoV-2 (pools S1-31) and SARS-CoV (pools S32-62) in duplicates. Sera of pooled healthy donors (n=13) were carried out in parallel. Data were normalized against healthy controls and the subtracted OD values are presented in a heat-map. Blue and pink indicate low and high OD values, respectively. **(d-e)** Determination of SARS-CoV-2 specific and pan-CoV linear B cell epitopes on S protein. **(d)** Sera of COVID-19 patients (n=6) were subjected to peptide-based ELISA for IgG detection using individual peptides of SARS-CoV-2 S peptide pools 14 and 21. The z-score values of patient were calculated using the formula [value – average (OD value of patient) / standard deviation (OD value of patient)]. Data shown are from two independent experiments and presented as mean. **(e)** Peptide binding response of COVID-19 patients on SARS-CoV-2 peptides S14P5 and S21P2, and the corresponding regions on SARS-CoV peptides S45P3 and S51P5, respectively. Statistical analysis was carried out with paired parametric two-tailed t-test (\**p*<0.05).

Next, we assessed the antigenic targets of these sera using a linear B-cell peptide library spanning the entire S protein of either SARS-CoV or SARS-CoV-2 with pools of five overlapping peptides (Figure 1c, Supplementary Figure 1b). Interestingly, two distinct peptide pools from SARS-CoV-2 S library, pools S14 and S21, were strongly detected by sera from COVID-19 patients, and not by recovered SARS patients (17 years post disease recovery) (Figure 1c, Supplementary Figure 1b). Sera from recalled SARS patients could neutralize SARS-CoV, but not the SARS-CoV-2 pseudotyped lentiviruses (Supplementary Figure 1c). Moreover, COVID-19 patients sera could strongly detect SARS-CoV S library pool S51, which partially overlaps with SARS-CoV-2 pool S21 (Figure 1c, Supplementary Figure 1b). This region encompasses the fusion peptide, which is highly conserved among coronaviruses [10, 11], suggesting a potential pan-coronavirus epitope at this location. Interestingly, targeting this region was also demonstrated to neutralize infection with a pan-coronavirus fusion inhibitor peptide [12]. Surprisingly, no linear epitope was identified in the receptor binding domain (RBD) which suggest that antibodies targeting that region are mostly conformational epitopes. Further assessment of individual peptides within pools S14 and S21 narrowed down the specific region of interest to peptides S14P5 and S21P2, respectively (Figure 1d). Recognition of these regions was stronger for the peptides of SARS-CoV-2 than SARS-CoV (Figure 1e). Together, these findings suggest that these linear B-cell epitopes are dominant antigenic regions, which corroborated previous bioinformatics predictions [7].

Using a recently published structure of SARS-CoV-2 S protein in prefusion conformation [2], peptide S14P5 is localized in proximity to the RBD (Figure 2a). As such, it is plausible that antibodies binding to this region may sterically hinder binding to ACE2 receptor, thereby abolishing virus infection [13]. Another possibility could be an allosteric effect on ACE2 binding. Similarly, peptide S21P2 contains a part of the fusion peptide sequence, which may potentially affect virus fusion (Figure 2b). In order to assess the importance of these regions in controlling SARS-CoV-2 infection, antibody depletion assays were performed against S14P5 and S21P2 (Figure 2c). Depletion efficiency and specificity was validated by ELISA, showing that only antibodies against the respective peptides were depleted but not other unrelated antibodies (Figure 2d). Interestingly, sera that were depleted for antibodies targeting either peptides S14P5, S21P2, or S14P5+S21P2 led to a significantly reduced ability to neutralize SARS-CoV-2 pseudovirus infection, as compared to the non-depleted sera controls (Figure 2e).

**Figure 2.**
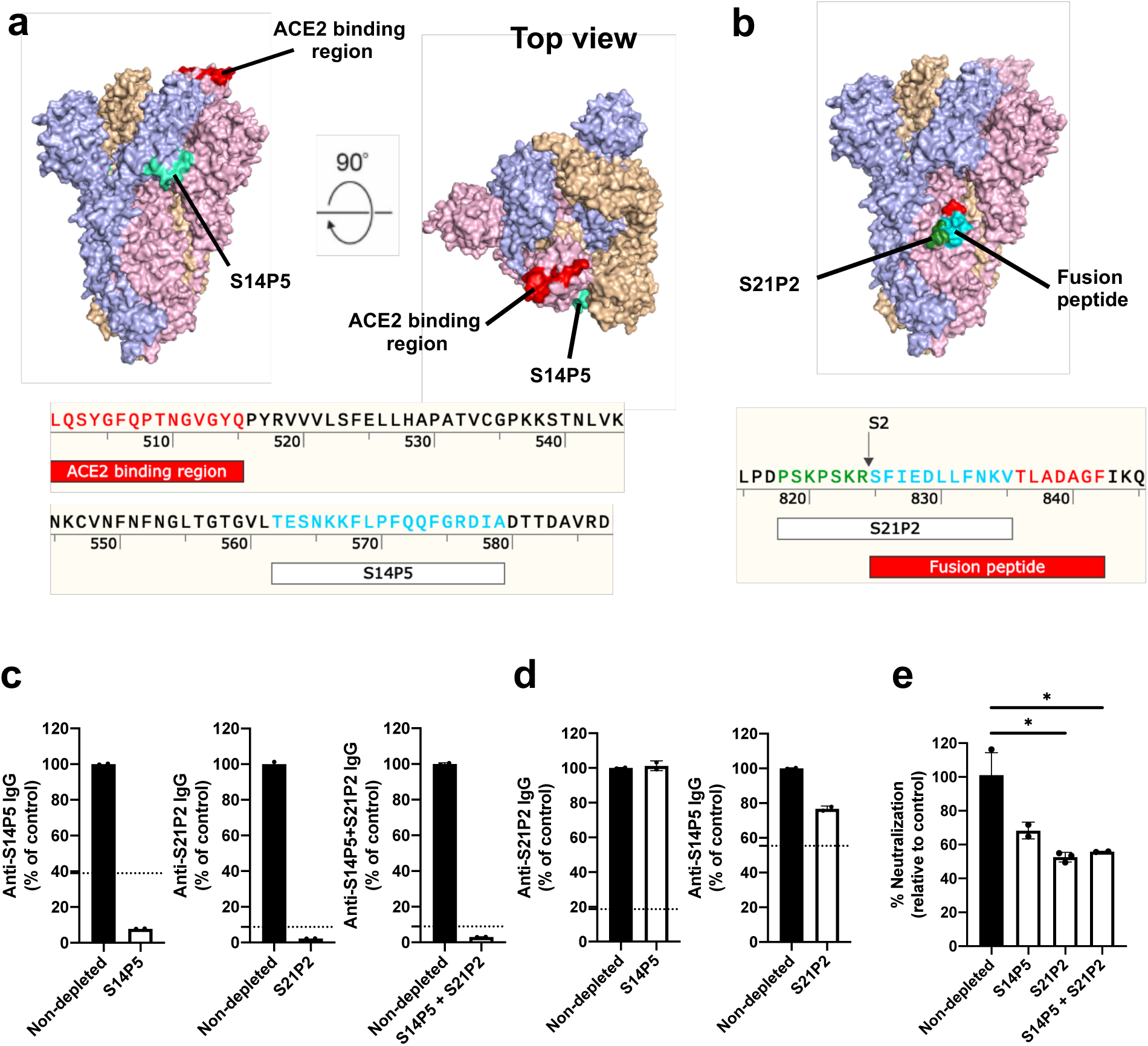
Antibodies against S14P5 and S21P2 linear B-cell epitopes neutralize SARS-CoV-2. **(a-b)** Localization and sequences of **(a)** SAR-CoV-2 specific S14P5 and **(b)** pan-CoV S21P2 epitopes on spike (S) protein (PDB: 6VSB) are shown. Each S monomer is denoted as either pink, blue or orange. **(c-e)** Pooled sera of COVID-19 patients (n=6) were added to plates coated with the corresponding peptides to deplete specific antibodies. **(c)** Non-depleted and peptide-specific antibody-depleted sera were then subjected to peptide-based ELISA of the peptide **(d)** or were cross-assayed. Data of depleted sera (white) were normalized to percentages of the non-depleted sera (black). Signal from pooled healthy donor sera were displayed as black dotted lines and data are shown as mean ± SD. **(e)** Non-depleted and peptide-specific antibody-depleted pooled sera were then mixed with SARS-CoV-2 pseudovirus before incubating with CHO-ACE2 cells for 48 h. The percentage of neutralization against pseudovirus infection, relative to the non-depleted sera, are shown. Data are presented as mean ± SD, and signal from pooled healthy donor sera were displayed as black dotted lines. Statistical analysis was carried out with one-sample t test (\**p*<0.05, \*\**p*<0.01).

Our results demonstrate that the two B-cell linear epitopes identified in this study are immunodominant. Depletion assays functionally validated that antibodies targeting S14P5 and S21P2 peptides possess significant neutralizing roles against SARS-CoV-2 pseudotyped lentiviruses. Importantly, we also assessed the potential presence of mutations in the peptide regions and found a low rate of mutations for S14P5 and S21P2 with low to moderate impact on the viral sequence (2 and 24 mutations out of 2596 viral genome sequences respectively, supplementary table 3) [14]. These results are essential to guide the design and evaluation of efficient and specific serological assays, as well as help prioritize vaccine target designs during this unprecedented crisis.

## Acknowledgements

Authors would like to thank Professor Yee-Joo Tan (Department of Microbiology, NUS) who kindly provided CHO-ACE2 cells and pXJ3’-S plasmid, and Assoc. Prof. Brendon Hanson (Defence Science Organization National Laboratories) who kindly provided pTT5LnX-CoV-SP plasmid. We would also like to thank the study participants who donated their blood samples to this project, and the healthcare workers who are caring for patients with COVID-19.

## Author contributions

CMP, GC, SNA, CYPL conceptualized, designed, acquired, analyzed, interpreted the data and wrote the manuscript. BW acquired, analyzed, interpreted the data and wrote the manuscript. RSLC, WHL and NKWY acquired and analyzed the data. YSL, MICC, SYT, LC, SK, SYT, BEY, and DCL designed and supervised sample collection. CIW, LR, LFPN conceptualized, designed, analyzed and wrote the manuscript. All authors revised and approved the final version of the manuscript.

