## Supplementary material for "Potent neutralizing antibodies in the sera of convalescent COVID-19 patients are directed against conserved linear epitopes on the SARS-CoV-2 spike protein": Supp methods

### Supplemental Information

#### Methods

#### Ethics statement

Written informed consent was obtained from participants in accordance with the tenets of the Declaration of Helsinki. For COVID-19 serum collection “A Multi-centred Prospective Study to Detect Novel Pathogens and Characterize Emerging Infections (The PROTECT study group)”, a domain specific review board (DSRB) evaluated the study design and protocol, which was approved under study number 2012/00917. Serum collection of SARS recall “Comparison of host immune responses to coronavirus infections” (ref) was approved by DSRB under study number 2020/00091. Sera from healthy volunteers “Study of blood cell subsets and their products in models of infection, inflammation and immune regulation” was approved under study number 2017/2512.

#### Patient Serum

Serum was collected in BD Vacutainer SST II Advance tubes (Fisher Scientific, #12927696). After clotting, serum was separated using centrifugation for 10 minutes at 1000 rcf, and aliquoted before storing at -80 °C. Patient serum was heat-inactivated for 30 minutes at 56°C before usage for this study. Patient information is provided in table 1.

Table 1: Details of patients individually tested for the library screen

| Patient ID | ICU | Sex | Age | Serum collection time (days post disease onset) |
|---|---|---|---|---|
| 2 | no | M | 35 | 17 |
| 4 | no | M | 41 | 15 |

| | | | | |
|---|---|---|---|---|
| 6 | no | F | 47 | 15 |
| 7 | no | F | 53 | 27 |
| 11 | no | M | 51 | 23 |
| 20 | yes | M | 56 | 30 |

**Linear peptide library**

The sequence used for the design of biotinylated linear peptides of the S glycoprotein of SARS-CoV and SARS-CoV-2 are under GenBank accession numbers NC_004718.3 and MN908947.3. Preliminary epitope screening was used with a library of peptides (Mimotopes, Mulgrave, VIC, Australia) consisting of 18-mer overlapping sequences. Peptides were used individually or as pooled sets. Five peptides were combined to form one pooled peptide set. Lyophilized individual peptides were dissolved in 200 µL of DMSO (Sigma-Aldrich, #D8418-100ML) to obtain a stock solution.

**Peptide-based ELISA**

B-cell linear library ELISA was performed as previously described (*1*). Briefly, streptavidin-coated plates (Thermo Fisher Scientific, #15125) were blocked with 0.1% PBST (0.1% v/v Tween-20, Sigma-Aldrich, #P1379-500ML, in PBS, Gibco, #20012-043) containing 1% w/v sodium caseinate (Sigma-Aldrich, #C8654-500G, lot BCBP6469) and 1% w/v bovine serum albumin (BSA; Sigma-Aldrich, #A7030-500G, lot SLBW5033) overnight at 4°C, before addition of pooled biotinylated peptides (1:1000 dilution in 0.1% PBST), followed by heat-inactivated patient serum samples (1:1000 dilution in 0.1% PBST). HRP-conjugated goat anti-human IgG (H+L) antibody (Jackson ImmunoResearch, # 109-035-088, lot 139159) prepared in 0.1% blocking buffer was used for detection of peptide-bound antibodies. 100 µL of TMB substrate

(Sigma-Aldrich, # T8665, lot SLCB5343) was used for a 5 min development and was stopped by addition of 100 µL of 0.16 M sulfuric acid prepared from 95-97% Sulfuric Acid stock solution (Merck, #1.00731.1000), prior to absorbance measurements. Absorbance was measured with the following parameters: 450 nm minus 690 nm (bandwidth of 9nm) in 5 flashes after a 10 sec shaking at 1 mm amplitude on an Infinite M200 plate reader (Tecan, firmware V_2.02_11/06).

**Peptide-specific antibody depletion of pooled SARS-CoV-2 sera**

Peptide-specific antibody depletion was performed as previously described (*2*) with some slight modifications. Briefly, selected synthetic biotinylated peptides were added at 1:1000 dilution in 0.1% PBST to pre-blocked streptavidin-coated plates and incubated at room temperature for 1 hour. Plates were washed 3 times with 0.1% PBST followed by PBS wash to remove traces of tween. Pooled patient sera were prepared at a dilution of 1:100 in DMEM (HyClone, #SH30243.01, lot AE29431634), and 50 µl per well of samples were added and incubated for 20 mins at room temperature for adsorption. The unbound portion was collected after 24 rounds of adsorption. ELISA analysis was performed as described before but at 1:2000 dilution, to assess the levels of selected antibodies before and after affinity depletion. Adsorbed samples were then mixed with lentiviruses pseudotyped with SARS-CoV-2 S protein as described below. Selected peptide sequences are given in table 2.

Table 2: List of individual peptide sequences tested

Will be in published version

**Cell lines and cell culture**

The human embryonic kidney epithelial cell 293T (ATCC, CRL-3216) was cultured in Dulbecco's modified Eagle's medium (Hyclone, SH30022.01) supplemented with 10% heat-inactivated FBS (Gibco, 10270-106). A stable cell line expressing human ACE2, CHO-ACE2 (a kind gift from Professor Yee-Joo Tan, Department of Microbiology, NUS, Singapore)(*3*) was maintained in Dulbecco's modified Eagle's medium supplemented with 10% heat-inactivated FBS, 1% MEM Non-Essential Amino Acids Solution (Gibco, 11140-050) and 0.5mg/ml of Geneticin™ Selective Antibiotic (Gibco, 10131-027). Every 2-3 days, cells were passaged by dissociating the cells with StemPro™ Accutase™ Cell Dissociation Reagent (Gibco, A1110501). ACE2 surface expression on CHO-ACE2 cells was verified using anti-human ACE2 AF647 (Santa Cruz Biotech, sc-390851, lot B0320).

**Generation of pseudotyped viral particles expressing SARS-CoV-2 S glycoprotein**

Based on the third-generation lentivirus system, pseudotyped viral particles expressing SARS-CoV-2 S protein was produced by reverse transfection of 30 x $10^6$ of 293T cells with 12µg pMDLg/pRRE (Addgene #12251), 6µg pRSV-Rev (Addgene #12253), 12µg pTT5LnX-coV-SP (a kind gift from DSO National Laboratories) and 24µg pHIV-Luc-ZsGreen (Addgene #39196) using Lipofectamine 2000 transfection reagent (Invitrogen, #11668-019) and cultured at 37°C incubator for 3 days. Viral supernatant was harvested, spun down by centrifugation to remove cell debris and filtered through a 0.45µm filter unit (Sartorius, #16555). Lenti-X p24 rapid titer kit (Takara Bio, #632200) was used to quantify the viral titres following the manufacturer instructions.

**Neutralization assay against SARS-CoV-2 S glycoprotein pseudotyped lentiviruses**

CHO-ACE2 cells were seeded at a density of 2.5 x $10^4$ cells in 100 μL of complete medium without Geneticin in 96-well Flat Clear Bottom Black Polystyrene TC-treated Microplates (Corning, #3904). After heat-inactivation at 56°C for 30 minutes, serially diluted patient serum were incubated in a 96-well flat-bottom cell culture plate (Costar, #3596) with an equal volume of pseudotyped virus (12ng of p24) at the final volume of 50μL at 37°C for 1 hour, and the mixture was added to the monolayer of pre-seeded CHO-ACE2 cells. After 1 hour of pseudotyped viral infection at 37°C, 150μl of culture medium was added to each well and the cells were further incubated for another 48 hours. Upon removal of culture medium, cells were washed twice with sterile PBS, and then lysed in 20μL of 1x Passive lysis buffer (Promega, E1941) with gentle shaking at 400rpm at 37°C for 30 minutes. Luciferase activity was then assessed using Luciferase Assay System (Promega, E1510) on a Promega GloMax Luminometer. $IC_{50}$ titer is the reciprocal of sera dilution at which half-maximal neutralization was calculated using GraphPad Prism (version 8.0.0).

**Data visualization and statistical analysis**

Structural data of SARS-CoV-2 S protein was retrieved from Protein Databank (PDB ID 6VSB) in homotrimeric prefusion conformation and visualized using PyMOL (Schrodinger, version 2.2.0). Heat-maps were generated using GraphPad Prism (version 8.0.0.).

Table 3: List of mutations that appear in from a list of 2596 sequences that were curated by China National Center for Bioinformation (CNCB) extracted on 30 March 2020 https://bigd.big.ac.cn/ncov (*4*)

| Peptide | Genome position | Virus number with variation | Annotation Type | Protein.Position.Amino acids change | Gene.Position.Codons | Impact Ensembl Variation - Calculated variant consequences"> | Last Update |
|---|---|---|---|---|---|---|---|
| | 23202 | 1 | missense_variant | QHD43416.1:p.547T>I | gene-S:c.1640aCa>aTa | MODERATE | 2020-03-29 20:27:38 |
| S14P5 | 23242 | 1 | synonymous_variant | QHD43416.1:p.560L | gene-S:c.1680ctG>ctT | LOW | 2020-03-29 20:27:38 |
| | 23347 | 1 | synonymous_variant | QHD43416.1:p.595V | gene-S:c.1785gtC>gtT | LOW | 2020-03-29 20:27:38 |
| | 23403 | 717 | coding_sequence_variant; missense_variant | QHD43416.1:p.614-; QHD43416.1:p.614D>G | gene-S:c.1841gAt>gRt; gene-S:c.1841gAt>gGt | MODIFIER; MODERATE | 2020-03-29 20:27:38 |
| | 23986 | 1 | synonymous_variant | QHD43416.1:p.808D | gene-S:c.2424gaT>gaC | LOW | 2020-03-29 20:27:38 |
| S21P2 | 23987 | 2 | missense_variant | QHD43416.1:p.809P>S | gene-S:c.2425Cca>Tca | MODERATE | 2020-03-29 20:27:38 |
| | 23989 | 1 | synonymous_variant | QHD43416.1:p.809P | gene-S:c.2427ccA>ccG | LOW | 2020-03-29 20:27:38 |
| | 23996 | 1 | missense_variant | QHD43416.1:p.812P>S | gene-S:c.2434Cca>Tca | MODERATE | 2020-03-29 20:27:38 |
| | 24011 | 1 | missense_variant | QHD43416.1:p.817F>L | gene-S:c.2449Ttt>Ctt | MODERATE | 2020-03-29 20:27:38 |
| | 24023 | 2 | missense_variant; synonymous_variant | QHD43416.1:p.821L>I; QHD43416.1:p.821L | gene-S:c.2461Cta>Ata; gene-S:c.2461Cta>Tta | MODERATE; LOW | 2020-03-29 20:27:38 |
| | 24034 | 17 | synonymous_variant; coding_sequence_variant | QHD43416.1:p.824N; QHD43416.1:p.824- | gene-S:c.2472aaC>aaT; gene-S:c.2472aaC>aaY | LOW; MODIFIER | 2020-03-29 20:27:38 |
| | 24054 | 9 | missense_variant | QHD43416.1:p.831A>V | gene-S:c.2492gCt>gTt | MODERATE | 2020-03-29 20:27:38 |
