## Supplementary material for "Potent neutralizing antibodies in the sera of convalescent COVID-19 patients are directed against conserved linear epitopes on the SARS-CoV-2 spike protein": Supp Figure 1

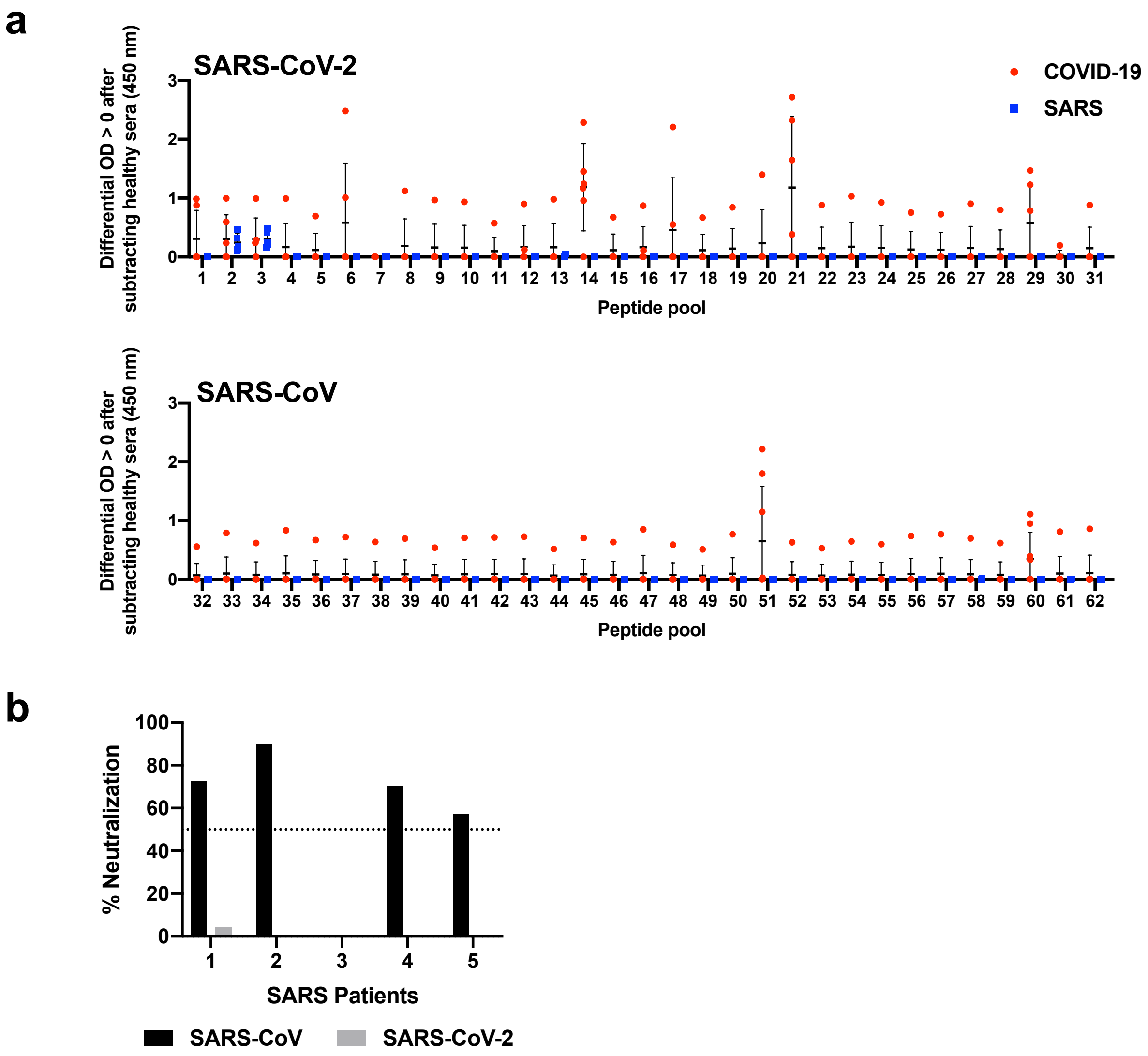


**Supplementary Figure 1. Antibody responses of COVID-19 and recovered SARS patients. (a)** Preliminary mapping of linear B-cell epitopes within SARS-CoV-2 and SARS-CoV spike (S) protein. Sera of COVID-19 (n=6) and SARS patients (n=5) at 1:1000 dilution were subjected to peptide-based ELISA for IgG using peptide pools from the S protein of SARS-CoV-2 (pools S1-31) and SARS-CoV (pools S32-62) in duplicates. Sera of pooled healthy donors (n=13) were carried out in parallel. Subtracted values are presented after normalization against healthy controls. Data is shown as mean ± SD. **(b)** Sera of recovered SARS patients (n=5; 1:100 dilution) were mixed with pseudotyped lentiviruses expressing SARS-CoV or SARS-CoV-2 S glycoprotein protein, prior to incubation with CHO-ACE2 cells for 48 h. Infection levels were determined by luciferase assay, and percentage neutralization was analyzed. Dotted line indicates 50% neutralization efficiency.

**Supp. Figure 1 Poh_Carissimo_Wang *et al*, 2020**
